# SARS-CoV-2 501Y.V2 escapes neutralization by South African COVID-19 donor plasma

**DOI:** 10.1101/2021.01.18.427166

**Authors:** Constantinos Kurt Wibmer, Frances Ayres, Tandile Hermanus, Mashudu Madzivhandila, Prudence Kgagudi, Brent Oosthuysen, Bronwen E. Lambson, Tulio de Oliveira, Marion Vermeulen, Karin van der Berg, Theresa Rossouw, Michael Boswell, Veronica Ueckermann, Susan Meiring, Anne von Gottberg, Cheryl Cohen, Lynn Morris, Jinal N. Bhiman, Penny L. Moore

## Abstract

SARS-CoV-2 501Y.V2 (B.1.351), a novel lineage of coronavirus causing COVID-19, contains substitutions in two immunodominant domains of the spike protein. Here, we show that pseudovirus expressing 501Y.V2 spike protein completely escapes three classes of therapeutically relevant antibodies. This pseudovirus also exhibits substantial to complete escape from neutralization, but not binding, by convalescent plasma. These data highlight the prospect of reinfection with antigenically distinct variants and foreshadows reduced efficacy of spike-based vaccines.

## Article

Individuals infected with severe acute respiratory syndrome coronavirus 2 (SARS-CoV-2), the virus that causes coronavirus disease 2019 (COVID-19), develop neutralizing antibodies that can persist for months^1,2^. Neutralizing antibodies are considered the primary correlate of protection from infection and are being pursued as therapeutics^3,4^. Interim analyses with monoclonal neutralizing antibodies have shown success, facilitating their authorization for emergency use^5,6^.

The SARS-CoV-2 receptor binding domain (RBD) exists in either an ‘up’ (receptor-accessible) or ‘down’ (receptor-shielded) conformation. RBD is the dominant neutralization target for this and other human coronaviruses^7,8^. These antibodies can be broadly divided into four main classes, of which two overlap with the angiotensin converting enzyme 2 (ACE2) receptor binding site (Fig. 1a and Supplementary Fig. 1a)^9^. Class 1 antibodies are most frequently elicited in SARS-CoV-2 infection and include a public antibody response to an epitope only accessible in the RBD ‘up’ conformation^10^. Class 2 antibodies use more diverse VH-genes and bind to RBD ‘up’ and RBD ‘down’ conformations of spike. After RBD, the N-terminal domain (NTD) of spike is the next most frequently targeted by neutralizing antibodies, most of which target a single immunodominant site^11^.

**Fig. 1.**
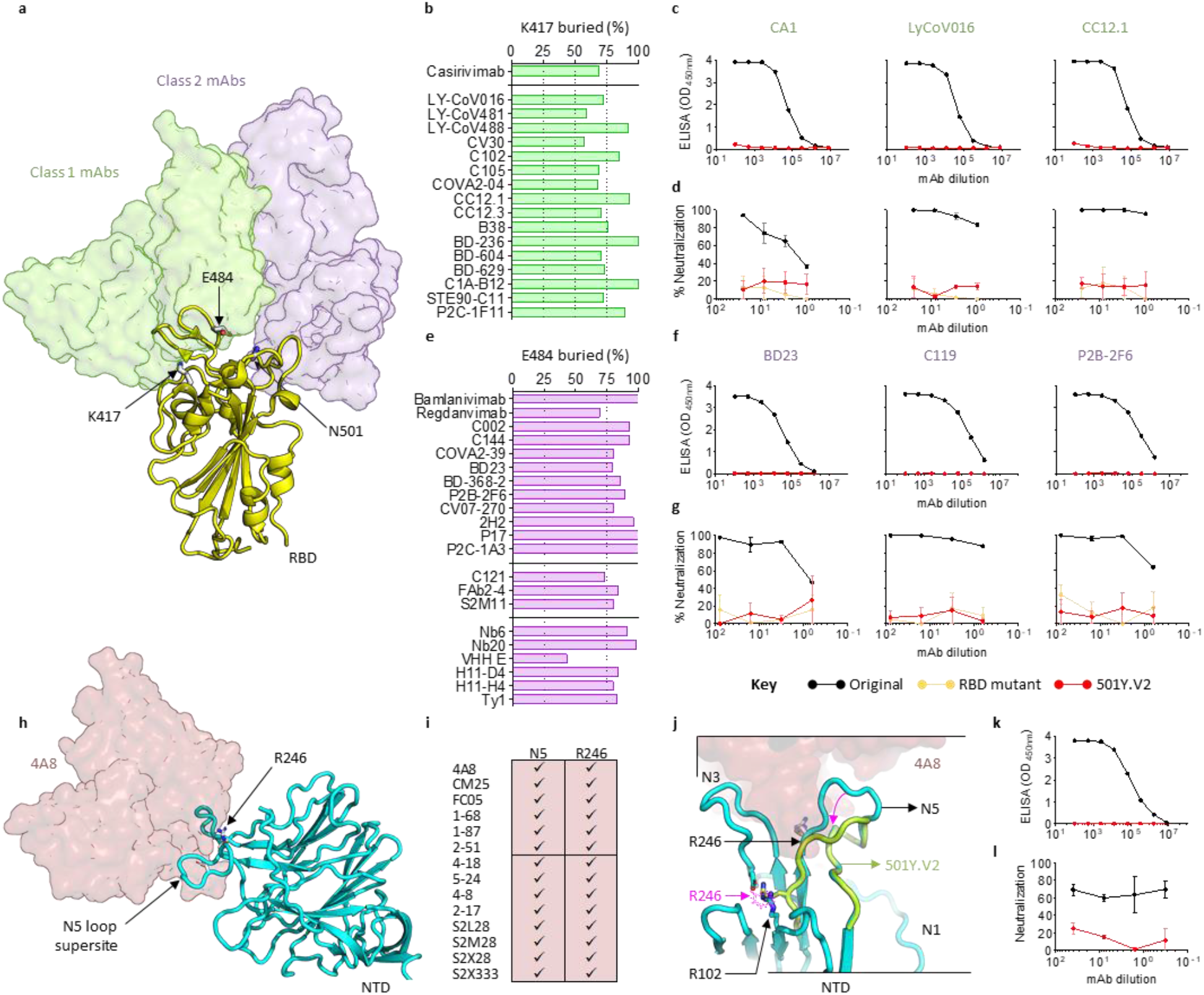
SARS-CoV-2 501Y.V2 is resistant to monoclonal antibodies. **a**, Structure of SARS-CoV-2 RBD (yellow) modeled in complex with class 1 (translucent green) or class 2 (translucent purple) neutralizing antibodies. Side chains of residues K417, E484 and N501 are indicated. mAb, monoclonal antibody. **b**, A plot showing percentage of K417 accessible surface area (x axis) buried (buried surface area) in class 1 antibody paratopes (listed on the y axis). VH3-53/66 antibodies are separated below the horizontal line. **c**, ELISA binding for CA1, LyCoV016 and CC12.1 to the original (black) or the 501Y.V2 RBD (red). **d**, Neutralization curves for the same antibodies shown in c, against the original pseudovirus (black), 501Y.V2 (red) or a chimeric construct that includes only the RBD substitutions K417N, E484K and N501Y (orange). **e**, Percentage of E484 accessible surface area buried in class 2 antibody paratopes (listed on y axis). VH1-2 antibodies (middle) or sy-/nanobodies (bottom) are separated with horizontal lines. **f**, ELISA binding for BD23, C119 and P2B-2F6 to the original (black) or 501Y.V2 RBD (red). **g**, Neutralization curves for the same antibodies shown in f, against original (black), 501Y.V2 (red) or RBD chimeric pseudoviruses (orange). **h**, Structure of SARS-CoV-2 NTD (cyan) modeled in complex with VH1-24 neutralizing antibody (translucent maroon). The N5-loop supersite and residue R246 are indicated. **i**, Contribution of N5 loop and R246 to NTD-directed neutralizing antibodies is indicated. **j**, Modeling of the Δ242-244 deletion (lime green). NTD loops N1, N3 and N5 are shown and the position of R246 in the original NTD and 501Y.V2 NTD is labeled with black and pink, respectively. The minimum displacement for 501Y.V2 loop N5 and the accompanying clash with R102 are indicated with pink arrows. **k**, ELISA binding for 4A8 to original (black) or 501Y.V2 NTD (red). **l**, Neutralization curves for 4A8 against the original (black) or 501Y.V2 (red) pseudovirus. All experiments were performed in duplicate.

We, and others, recently described a new SARS-CoV-2 lineage in South Africa, defined as Nextstrain clade 20H/501Y.V2 (PANGOLin lineage B.1.351)^12^. This lineage is defined by nine changes in the spike protein (Supplementary Fig. 1b) relative to the Wuhan-1 D614G spike mutant that previously dominated in South Africa (here referred to as the original lineage)^13^. These changes include N501Y, which confers enhanced affinity for ACE2^14^, and clusters of substitutions in two immunodominant regions of spike, suggesting escape from neutralization. Indeed, substitutions at E484 reduce neutralization sensitivity to convalescent plasma^15^. We therefore compared neutralization by monoclonal antibodies and convalescent plasma of 501Y.V2 to Wuhan-1 D614G, using a spike-pseudotyped lentivirus neutralization assay.

An analysis of 17 class I antibody structures revealed their epitopes to be centered on spike residue K417, one of three substitutions in the RBD of the 501Y.V2 lineage. These antibodies contact 60–100% of residue K417 side-chain-accessible surface area, including key hydrogen bonds at this site (Fig. 1b). Three representative antibodies were assessed by ELISA and achieved saturated binding to recombinant RBD from the original lineage but not 501Y.V2 RBD (Fig. 1c). Similarly, all three antibodies potently neutralized the original lineage, but not the 501Y.V2 pseudovirus (at 25 μg.ml^−1^), confirming dependence on the K417 residue (Fig. 1d).

A structural analysis of 15 class 2 antibodies and 6 nanobodies revealed key interactions with spike residue E484 (Fig. 1e). Each contacted 40–100% of the E484 side-chain-accessible surface area and formed critical hydrogen bonds or charged interactions at this site. As with class 1 antibodies, three representative class 2 antibodies failed to bind 501Y.V2 RBD (Fig. 1f) and were unable to neutralize the 501Y.V2 pseudovirus (Fig. 1g). Thus, the SARS-CoV-2 501Y.V2 lineage has effectively escaped two major classes of neutralizing antibodies targeting an immunodominant, highly antigenic site in the RBD of the spike protein.

501Y.V2 is also defined by several changes in the NTD, including a three-amino-acid deletion preceding the N5-loop supersite (Fig. 1h and Supplementary Fig. 1b). An analysis of NTD-bound antibody structures showed that roughly half their neutralization interfaces with spike comprised the N5-loop supersite, often involving key residue R246 (Fig. 1i). Modeling the 501Y.V2 N5-loop deletion onto the antibody 4A8-bound spike structure revealed an apical loop displacement of at least 8 Å away from the 4A8 paratope (Fig. 1j). In addition, the deletion would shift position R246 three amino acids earlier, bringing it into proximity with R102 and creating a potential clash that could locally disrupt the N5-loop conformation and shift it even further from the 4A8 paratope. Consequently, when assessed by ELISA, 4A8 was unable to bind to recombinant 501Y.V2 NTD (Fig. 1k) and failed to neutralize the 501Y.V2 pseudovirus (Fig. 1l), showing escape from N5-loop targeted neutralizing antibodies.

We next sought to evaluate the effect of 501Y.V2 spike substitutions on polyclonal plasma/sera derived from individuals with PCR-confirmed SARS-CoV-2 infection, including individuals who were hospitalized with severe COVID-19. Samples were divided into two groups, half with higher titer neutralizing antibodies (22 of 44, 50% inhibitory dilution (ID_50_) > 400) and half with lower titers (22 of 44, 400 ≥ ID_50_ > 25) to the original SARS-CoV-2 D614G lineage (Fig. 2 and Supplementary Fig. 2a). Consistent with previous studies, when stratified by disease severity, convalescent individuals who reported mild-to-moderate disease developed substantially lower neutralizing antibody titers (average ID_50_ titer 488, n = 30) than severely ill individuals from the hospitalized cohorts (average ID_50_ titer 4,212, n = 14).

**Fig. 2.**
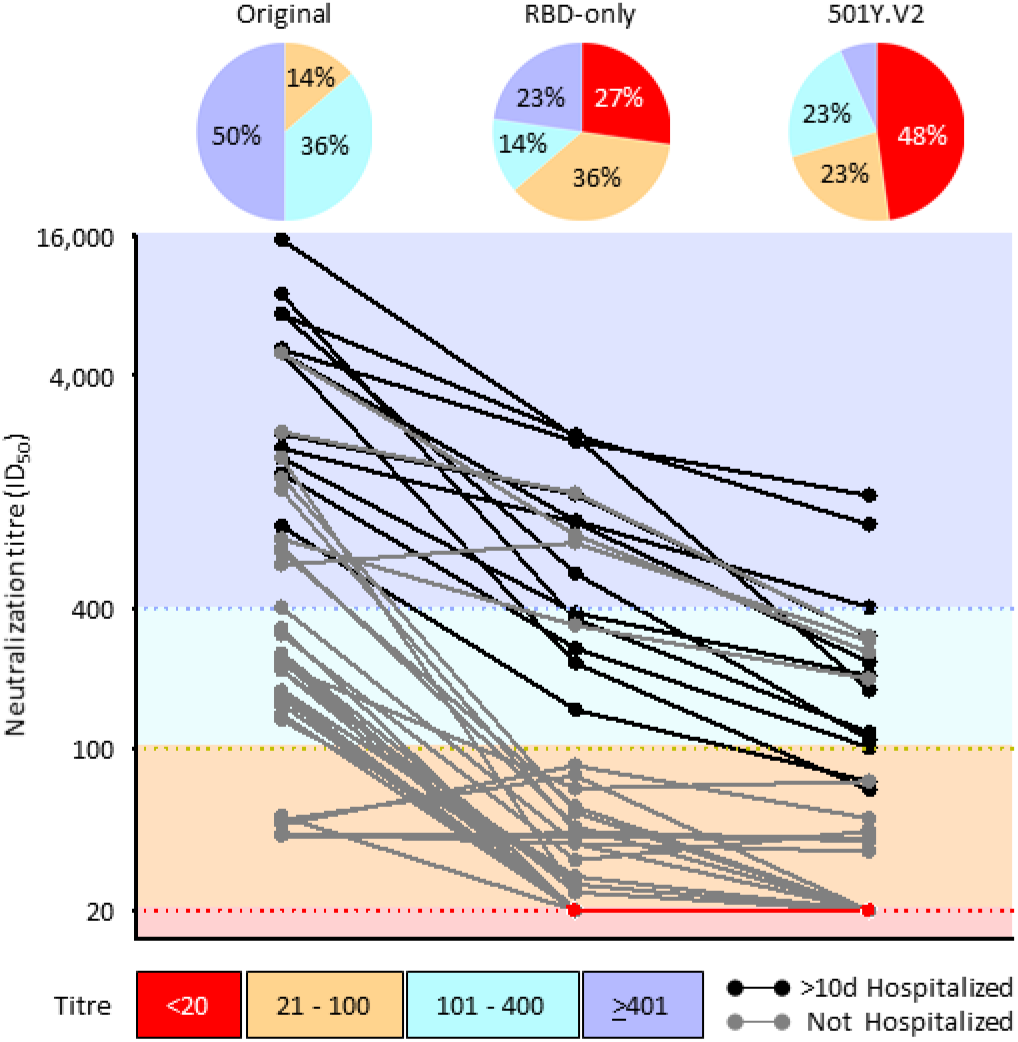
SARS-CoV-2 501Y.V2 increased resistance to neutralization by convalescent plasma/serum. Plasma/serum collected from individuals infected with SARS-CoV-2 was assessed for neutralization to the original lineage (Wuhan-1 D614G, left), an RBD chimeric mutant containing K417N, E484K and N501Y substitutions only (middle) or the 501Y.V2 lineage pseudovirus. Twelve of the samples were collected from donors hospitalized for >10 d with COVID-19 (black). The graph is colored according to the magnitude of neutralization titer, with ID_50_ greater or lesser than 1:400 colored dark or light blue, respectively and titer <100 colored orange. The limit of detection (knockout) was an ID_50_ < 20 (red). Pie charts above each set of data points summarize the proportion of samples in each titer group.

When these same samples were assessed against the 501Y.V2 pseudovirus, nearly half (21 of 44, 48%) had no detectable neutralization activity (and 71% had ID_50_ < 100). Only three samples (7%) retained titers of ID_50_ > 400 (Fig. 2 and Supplementary Fig. 2a). Notably, these three samples were obtained from individuals reporting severe disease and had among the highest neutralization titers against the original virus. Conversely, four samples with borderline neutralization of the original virus were unaffected by 501Y.V2 substitutions, perhaps representing additional, less-potent specificities. To define the location of dominant escape substitutions, neutralization was also assessed against the RBD chimeric pseudovirus containing only three 501Y.V2 substitutions (K417N, E484K and N501Y) (Fig. 2 and Supplementary Fig. 2a). Substantial loss of neutralization was also observed against the RBD-only mutant, with 27% of the samples losing all activity against the RBD triple mutants (63% had ID_50_ < 100) and only 23% retaining higher titers of ID_50_ > 400. These data provide more evidence for the dominance of class 1 and class 2 neutralizing antibodies in polyclonal sera; however, differences in neutralization between RBD-only chimera and 501Y.V2 also highlight the contribution of 501Y.V2 NTD substitutions (L18F, D80A, D215G and Δ242-244) to neutralization escape. This was particularly evident in higher titer samples, which retained an average ID_50_ titer of 680 against the RBD-only mutant.

While neutralizing antibodies to SARS-CoV-2 are dominated by the specificities defined above, non-neutralizing antibodies are also elicited during SARS-CoV-2 infection. To determine whether 501Y.V2 is still recognized by non-neutralizing antibodies, the binding of polyclonal sera (from Fig. 2) to a recombinant protein that includes the RBD and subdomain 1 (RBD + SBD1) of 501Y.V2 or the original lineage was assessed by ELISA (Supplementary Fig. 2b). These data revealed that binding of polyclonal plasma to 501Y.V2 RBD + SBD1 was only substantially affected in a minority of cases (14 of 44 with more than a fivefold reduction, 32%). Most of the convalescent plasma/serum suffered less than a four-fold reduction in total binding activity (as calculated by area under the curve), suggesting a considerable non-neutralizing antibody component is still able to bind the 501Y.V2 spike.

Among previous emerging lineages, only D614G has subsequently become globally dominant[16]. The repeated, independent evolution of spike position 501 in 501Y.V1 (https://virological.org/t/576), 501Y.V2^12^, and 501Y.V3 (https://virological.org/t/586), strongly argues for a selective advantage, likely enhanced transmissibility, of these new variants. Here we have shown that the 501Y.V2 lineage, containing nine spike substitutions, and rapidly emerging in South Africa during the second half of 2020, is resistant to neutralizing antibodies found in 48% of individuals infected with previously circulating lineages. These data, showing a 13-fold reduction in mean titre, are corroborated by vesicular stomatitis virus-pseudotyped and live virus assays showing an 11- to 33-fold and 6- to 204-fold reduction in mean titre (including complete knock out) relative to the original lineage, respectively^17,18^. The 501Y.V3 lineage has similar changes including 417T and 484K (in RBD) as well as 18F and 20N (in NTD), thus also having strong potential for high levels of neutralization resistance. The independent emergence and subsequent selection of 501Y lineages with key substitutions conferring neutralization resistance strongly argues for selection by neutralizing antibodies as the dominant driver for SARS-CoV-2 spike diversification and makes these lineages of considerable public health concern. This suggests that, despite the many people who have already been infected with SARS-CoV-2 globally and are presumed to have accumulated some level of immunity, new variants such as 501Y.V2 may pose a substantial reinfection risk.

While higher titers of neutralizing antibodies are common in hospitalized individuals, most people infected with SARS-CoV-2 develop low-to-moderate neutralization titers^2^. Therefore, the data herein suggest that most individuals infected with previous SARS-CoV-2 lineages will have greatly reduced neutralization activity against 501Y.V2. This dramatic effect on plasma neutralization can be explained by the dominance of RBD-directed neutralizing antibodies, supported by studies showing reduced plasma neutralization titers mediated by the E484K change alone^15^. Notably, here we show that the K417N change also has a crucial role in viral escape, effectively abrogating neutralization by a well-defined, multidonor class of VH3-53/66 germline-restricted public antibodies that comprise some of the most common and potent neutralizing antibodies to SARS-CoV-2^8^.

The marked loss of neutralization against 501Y.V2 pseudovirus compared to the RBD-only chimeric pseudovirus demonstrates the important role that substitutions in the NTD play in mediating immune escape. For 501Y.V2 this resistance to neutralization is likely mediated by a three-amino-acid deletion that completely disrupts a dominant public antibody response to the N5-loop supersite^11^. This deletion predominates among 501Y.V2 variants and occurs either alone or with an R246I substitution that is also important for neutralization by several NTD-directed neutralizing antibodies.

The relatively rapid acquisition of a comprehensive suite of neutralization escape substitutions likely occurred because of the large number of commonly shared public antibodies (such as VH3-53/66, VH1-2, and VH1-24) to both the RBD and NTD of spike, together with high levels of SARS-CoV-2 transmission. The sporadic emergence of escape substitutions in long-term viral shedders, including immunocompromised individuals, may also contribute to the emergence of neutralization-resistant viruses^19^. Altogether, these data highlight the need for increased, ongoing genomic surveillance during the SARS-CoV-2 pandemic.

Crucially, it is from these same public antibody responses that many therapeutic strategies currently under development have been derived^4^. The overwhelming majority of monoclonal antibodies already on the path to licensure target residues K417 or E484 and are therefore likely to be futile against 501Y.V2. In addition, emerging variants may limit the use of recently identified neutralizing antibodies that target the NTD N5-loop supersite. Some of these monoclonal antibodies have already been granted emergency use authorization in the United States (Regeneron Pharmaceuticals and Eli Lilly and Company), including antibodies ineffective against 501Y.V2 (such as REGN10933 and LY-CoV555) as well as antibodies likely to retain neutralization of this variant (REGN10987 and VIR-7831), some of which are being engineered to potentially enhance virus-specific T cell function (VIR-7832).

These data also have implications for the effectiveness of SARS-CoV-2 vaccines, largely based on immune responses to the original spike protein. Indeed, sera from the Moderna and Pfizer-BioNTech vaccinees show significantly reduced neutralization of 501Y.V2^18^. Furthermore, compared to global efficacy estimates, vaccine efficacy in South Africa (overlapping with the emergence of 501Y.V2) has been significantly reduced for several vaccines. Conversely, these same trials seem to retain efficacy against severe COVID-19. While antibody effector functions elicited by infection and vaccination have been implicated in protecting from reinfection and disease^20^, the role of non-neutralizing antibodies and the efficacy of T cell responses to 501Y.V2 remain to be elucidated. We did not measure the extent of these responses to 501Y.V2 but did show that a substantial proportion of non-neutralizing antibodies remain active against 501Y.V2 recombinant RBD protein. Ultimately, the correlates of protection against SARS-CoV-2 infection and severe COVID-19 disease remain undetermined and rely upon ongoing large-scale clinical trials. Nevertheless, these data highlight the urgent requirement for rapidly adaptable vaccine design platforms and the need to identify less-mutable viral targets for incorporation into future immunogens.

## Acknowledgements

We thank the COVID-19 convalescent plasma donors, the staff of the South African National Blood Services for contributing samples that enabled this work, Zelda van der Walt, Talita de Villiers, Wesley van Hougenhouck-Tulleken for contributing to patient management and sample collection and Amelia Buys for technical support. We thank the participants of the Novel Coronavirus (COVID-19) viral shedding and clinical characterization study (University of the Witwatersrand Health Research Ethics (Medical) committee reference M160667) who contributed samples and the GERMS-SA clinical staff for their contributions to sample and data collection. Rajiev Ramlall and Marthinus Heystek at Tshwane District Hospital are thanked for their input and assistance with facilitating participant recruitment for the Pretoria COVID-19 Study Cohort. We acknowledge funding from the South African Medical Research Council (Reference numbers 96825, SHIPNCD 76756 and DST/CON 0250/2012), The Wellcome Trust (Grant no 221003/Z/20/Z),) United States Centers for Disease Control and Prevention (Grant number 5 U01IP001048-05-00) and the ELMA South Africa Foundation (Grant number 20-ESA011). P.L.M. is supported by the South African Research Chairs Initiative of the Department of Science and Innovation and the National Research Foundation of South Africa (Grant No 98341). C.K.W. is supported by Fogarty International Center of the National Institutes of Health under Award Number R21TW011454 (This work is solely the responsibility of the authors and does not necessarily represent the official views of the National Institutes of Health) as well as the FLAIR Fellowship program under award number FLR\R1\201782 (The FLAIR Fellowship Programme is a partnership between the African Academy of Sciences and the Royal Society funded by the UK Government’s Global Challenges Research Fund). KvdB is supported in part by the Fogarty International Centre or the National Institutes of Health under Award Number 1D43TW010345. We thank Drs Nicole Doria-Rose, David Montefiori, Elise Landais and Michael Farzan for reagents and assistance in establishing the SARS-CoV-2 pseudotyped neutralization assay and enabling equivalency and proficiency testing. We thank Drs Devin Sok, Elise Landais, Dennis Burton, Nicole Doria-Rose and Peter Kwong for SARS-Co-V-2 directed monoclonal antibodies. We are grateful to Thandeka Moyo-Gwete and Zanele Molaudzi for expressing monoclonal antibodies. We thank the informal 501Y.V2 consortium of South African scientists, chaired by Drs Willem Hanekom and Tulio de Oliveira for suggestions and discussion of data. We also thank all NGS-SA laboratories in South Africa that were responsible for producing the SARS-CoV-2 genomes that enabled the rapid dissemination of SARS-CoV-2 sequences and the identification of 501Y.V2.

## Author contributions

CKW, JNB and PLM conceived the study, designed experiments, analysed data and wrote the paper. CKW, FA, BO, and BEL made molecular constructs, and expressed antibodies. CKW expressed and purified recombinant antigens. CKW, TH, MM, and PK made pseudoviruses, while TH, MM and PK performed and analysed neutralization experiments. JNB and TdO contributed to the identification of the 501Y.V2 lineage and the mutations comprised therein. LM contributed to establishing SARS-CoV-2 assays and infrastructure. MV, KvdB, TR, MB, VU, SM, AvG and CH provided sera from convalescent donors and associated clinical information. All authors critically reviewed and approved the final manuscript.

## Materials and Methods

### Samples and ethics approvals

Plasma/serum samples were obtained from individuals without human immunodeficiency virus (HIV), enrolled into one of three studies described below. All participants provided informed consent.

#### Hospitalized Steve Biko cohort

This study has been given ethics approval by the University of Pretoria, Human Research Ethics Committee (Medical) (247/2020). Serum samples were obtained (longitudinally) from hospitalized patients with PCR-confirmed SARS-CoV-2 infection, known HIV status and aged ≥18 years. Samples from six participants with symptom onset between May and August 2020 were used.

#### Novel coronavirus (COVID-19) viral shedding and clinical characterization study

This study has been given ethics approval by the University of the Witwatersrand Human Research Ethics Committee (Medical) M160667. Serum samples were obtained (longitudinally) from hospitalized patients with PCR-confirmed SARS-CoV-2 infection, known HIV status and aged ≥18 years. Samples from six participants with symptom onset between May and September 2020 were used.

#### South African National Blood Service

Plasma was obtained from blood donors of the South African National Blood Service (ethics clearance from South African National Blood Service Human Research Ethics Committee 2019/0519), who had PCR-confirmed SARS-CoV-2 infection, had recovered and were at least 28 d post-symptom onset. All donors met the standard eligibility criteria for donors who donate source plasma, which includes being generally healthy, being older than 18 years of age, weighing more than 55 kg and leading a lifestyle that reduces risk of acquiring transfusion transmissible infections. Only male and nulliparous females were accepted as COVID‐19 convalescent plasma donors.

### Structural modelling

The closed prefusion SARS-CoV-2 spike protein from Nextstrain^21^ clade 501Y.V2 (PANGOLin lineage B.1.351, (https://cov-lineages.org/pangolin.html) was modelled on PDBID 6VXX^22^, while a model of the RBD was averaged using neutralizing domain-bound structures compared to the ACE2-bound PDBID 6LZG^23^. Class 1, 2, 3 and 4 antibodies were represented with Fab domains from CV30 (6XE1), C104 (7k8u), VIR-7831 (6WPS) and EY6A (6ZCZ). The 501Y.V2 NTD domain was modeled using both antibody-constrained PDBID 7C2L^11^and unconstrained structures of spike deposited in the protein databank^24^. Involvement of the N5-loop supersite was estimated from available preprint data^25–29^. Buried interfaces were calculated using PDBePISA (https://www.ebi.ac.uk/pdbe/pisa/pistart.html) and analyzed according to neutralization class^30–33^. All structures were visualized in PyMol^34^.

### Expression and purification of SARS-CoV-2 antigens

Monoclonal antibodies or recombinant SARS-CoV-2 RBD + SBD1 or NTD proteins were transfected in FreeStyle 293F suspension cells (Life Technologies) using PEIMAX transfection reagent (Polysciences). Transfections were incubated at 220 r.p.m., 37 °C, 9% CO_2_ for 6–7 d and clarified supernatants were purified using nickel affinity or protein A and size-exclusion chromatography.

### ELISA

Recombinant RBD + SBD1 proteins were coated at 2 μg.ml^−1^ onto 96-well, high-binding plates and incubated overnight at 4 °C. The plates were then washed and incubated at 37 °C for 1–2 h in blocking buffer (5% skimmed milk powder, 0.05% Tween 20, 1× PBS). Plasma samples were added at 1:100 dilution and serially diluted fivefold in blocking buffer. Following a 1-h incubation at 37 °C, an anti-human horseradish peroxidase-conjugated antibody was added for 1 h at 37 °C. The signal was developed with TMB substrate (Thermo Fisher Scientific) for 5 min at room temperature, followed by addition of 1 M H_2_SO_4_ stop solution. Absorbance at 450 nm was measured and used to calculate area under the curve using GraphPad Prism v.8.4.2.

### Lentiviral pseudovirus production and neutralization assay

The 293T/ACE2.MF cells modified to overexpress human ACE2 were kindly provided by M. Farzan (Scripps Research). Cells were cultured in DMEM (Gibco BRL Life Technologies) containing 10% heat-inactivated serum (FBS) and 3 μg.ml^−1^ puromycin at 37 °C, 5% CO_2_. Cell monolayers were disrupted at confluency by treatment with 0.25% trypsin in 1 mM EDTA (Gibco BRL Life Technologies).

The SARS-CoV-2, Wuhan-1 spike, cloned into pCDNA3.1 was mutated using the QuikChange Lightning Site-Directed Mutagenesis kit (Agilent Technologies) to include D614G (original) or K417N, E484K, N501Y, D614G (RBD only) or L18F, D80A, D215G, Δ242-244, K417N, E484K, N501Y, D614G, A701V (501Y.V2). Pseudoviruses were produced by co-transfection with a lentiviral backbone (HIV-1 pNL4.luc encoding the firefly luciferase gene) and either of the SARS-CoV-2 spike plasmids with PEIMAX (Polysciences). Culture supernatants were clarified of cells by a 0.45 μM filter and stored at −70 °C.

Plasma/serum samples were heat-inactivated and clarified by centrifugation. Pseudovirus and serially diluted plasma/sera were incubated for 1 h at 37 °C, 5% CO2. Cells were added at 1 × 10^4^ cells per well after 72 h of incubation at 37 °C, 5% CO_2_, luminescence was measured using PerkinElmer Life Sciences Model Victor X luminometer. Neutralization was measured as described by a reduction in luciferase gene expression after single-round infection of 293T/ACE2.MF cells with spike-pseudotyped viruses. Titers were calculated as the reciprocal plasma dilution (ID_50_) causing 50% reduction of relative light units. Equivalency was established through participation in the SARS-CoV-2 Neutralizing Assay Concordance Survey Concordance Survey 1 run by EQAPOL and VQU, Duke Human Vaccine Institute. Cell-based neutralization assays using live virus or pseudovirus have demonstrated high concordance, with highly correlated 50% neutralization titers (Pearson *r* = 0.81–0.89)^35^.

## Statistical analyses

A Wilcoxon matched pairs signed-rank test showed significant difference in neutralization ID_50_ titer between the original pseudovirus and the RBD-only mutant or N501Y.V2, as well as between the RBD-only mutant and the 501Y.V2 lineage (*P* ≤ 0.0001).

## Data availability

Authors can confirm that all relevant data are included in the paper and/or its supplementary files.

## Supplementary information

**Supplementary Fig. 1.**
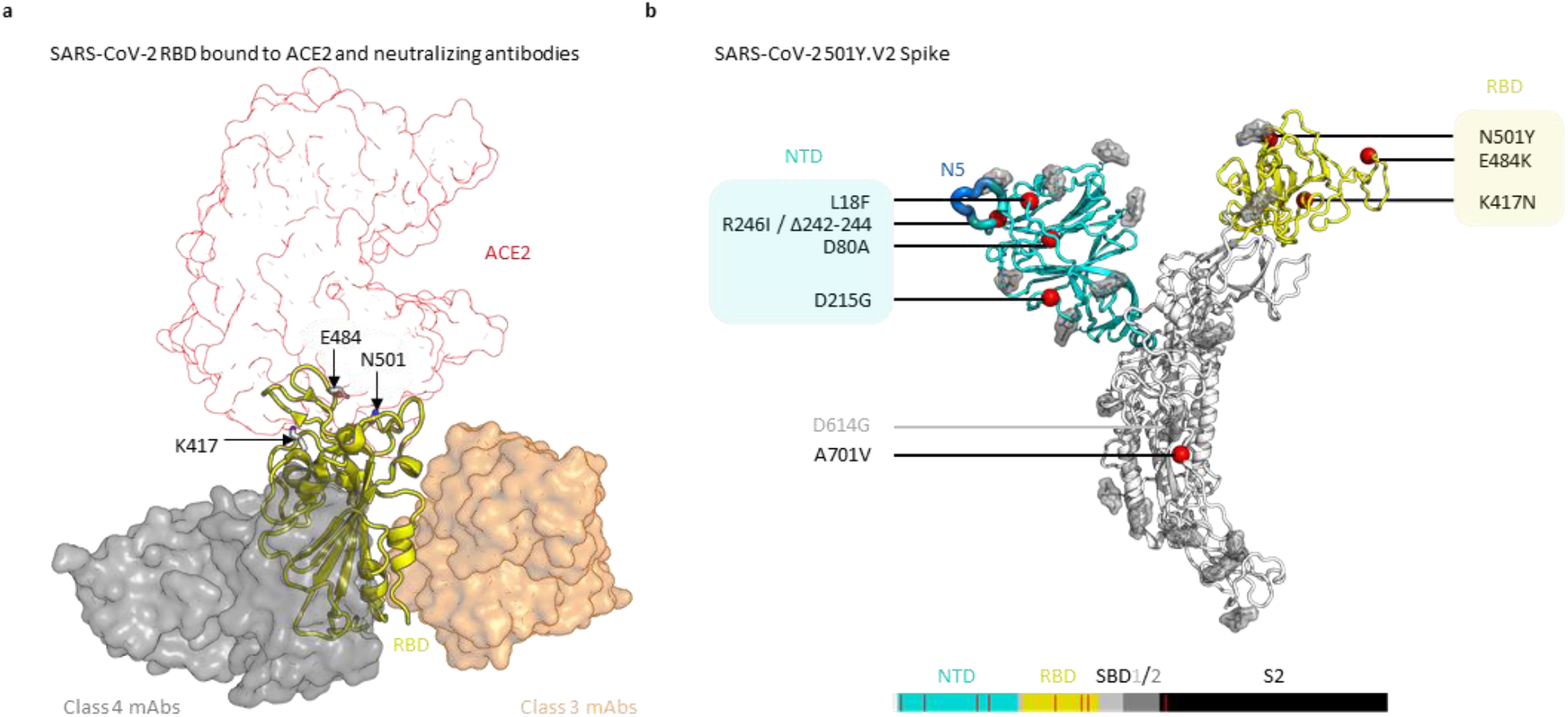
Location of 501Y.V2 defining mutations. **a**. The SARS-CoV-2 RBD is shown in yellow cartoon view, bound to human ACE2 (red outline, surface view), as well as a class 3 (translucent orange) or class 4 (translucent grey) neutralizing antibodies that are not affected by 501Y.V2 changes. The location of 501Y.V2 associated changes 417N, 484K, and 501Y are indicated. **b**. Cartoon schematic of a single SARS-CoV-2 spike protomer, with NTD and RBD coloured cyan and yellow, respectively. A single N-acetylglucosamine is shown at each glycosylation site, and the location of 501Y.V2 changes are shown with the red spheres and labelled. A linear schematic is also shown.

**Supplementary Fig. 2.**
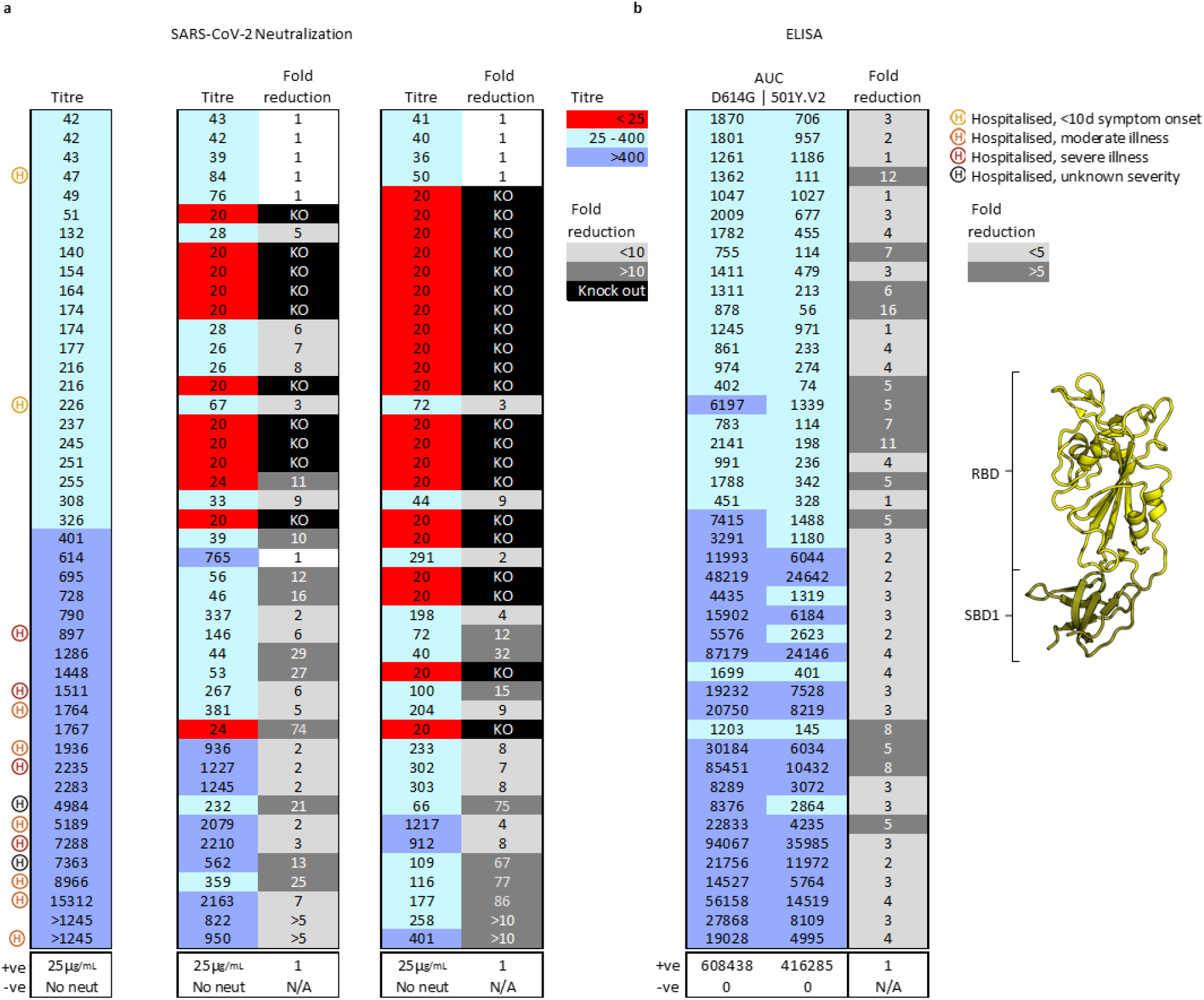
SARS-CoV-2 501Y.V2 shows increased resistance to neutralization but not binding by convalescent plasma/serum. **a**. Plasma/serum samples collected from SARS-CoV-2 infected individuals who were (n=14) or were not (n=30) hospitalized with COVID-19, ranked by titre against the original SARS-CoV-2 D614G lineage (column 1). Neutralization titre is coloured according to magnitude, where titres greater or lesser than 1:400 are coloured dark or light blue, respectively. Neutralization titre and fold decrease relative to the original lineage are shown for an RBD-only chimeric virus containing the K417N, E484K, and N501Y substitutions (columns 2 and 3), and the 501Y.V2 lineage virus (column 4 and 5). Neutralization titres <1:25 are coloured red, while a complete knock out of neutralization activity is highlighted in black. **b**. Binding of plasma samples (from Fig.2a) against the Receptor Binding Domain + Sub-Domain 1 (shown in yellow and olive cartoon view) from the original virus (column 1) or the 501Y.V2 lineage (column 2) and plotted as area under the curve. The fold reduction in AUC is shown in column 3. All experiments were performed in duplicate. In all instances, monoclonal antibodies CC12.23 or palivizumab were used as positive and negative controls, respectively.

## References

1. Zhou, P., et al. A pneumonia outbreak associated with a new coronavirus of probable bat origin. Nature 579, 270–273 (2020).

2. Wajnberg, A., et al. Robust neutralizing antibodies to SARS-CoV-2 infection persist for months. Science 370, 1227–1230 (2020).

3. Mercado, N.B., et al. Single-shot Ad26 vaccine protects against SARS-CoV-2 in rhesus macaques. Nature 586, 583–588 (2020).

4. Marovich, M., Mascola, J.R. & Cohen, M.S. Monoclonal Antibodies for Prevention and Treatment of COVID-19. JAMA 324, 131–132 (2020).

5. Weinreich, D.M., et al. REGN-COV2, a Neutralizing Antibody Cocktail, in Outpatients with Covid-19. N. Engl. J. Med. 384, 238–251 (2021).

6. Chen, P., et al. SARS-CoV-2 Neutralizing Antibody LY-CoV555 in Outpatients with Covid-19. N. Engl. J. Med. 384, 229–237 (2021).

7. Piccoli, L., et al. Mapping Neutralizing and Immunodominant Sites on the SARS-CoV-2 Spike Receptor-Binding Domain by Structure-Guided High-Resolution Serology. Cell 183, 1024–1042 e1021 (2020).

8. Barnes, C.O., et al. Structures of Human Antibodies Bound to SARS-CoV-2 Spike Reveal Common Epitopes and Recurrent Features of Antibodies. Cell 182, 828–842 e816 (2020).

9. Barnes, C.O., et al. SARS-CoV-2 neutralizing antibody structures inform therapeutic strategies. Nature 588, 682–687 (2020).

10. Yuan, M., et al. Structural basis of a shared antibody response to SARS-CoV-2. Science 369, 1119–1123 (2020).

11. Chi, X., et al. A neutralizing human antibody binds to the N-terminal domain of the Spike protein of SARS-CoV-2. Science 369, 650–655 (2020).

12. Tegally, H., et al. Emergence and rapid spread of a new severe acute respiratory syndrome-related coronavirus 2 (SARS-CoV-2) lineage with multiple spike mutations in South Africa. medRxiv (2020).

13. Tegally, H. et al. Sixteen novel lineages of SARS-CoV-2 in South Africa. Nat. Med. https://doi.org/10.1038/s41591-021-01255-3 (2021).

14. Starr, T.N., et al. Deep Mutational Scanning of SARS-CoV-2 Receptor Binding Domain Reveals Constraints on Folding and ACE2 Binding. Cell 182, 1295–1310 e1220 (2020).

15. Weisblum, Y., et al. Escape from neutralizing antibodies by SARS-CoV-2 spike protein variants. Elife 9 (2020).

16. Korber, B., et al. Tracking Changes in SARS-CoV-2 Spike: Evidence that D614G Increases Infectivity of the COVID-19 Virus. Cell 182, 812–827 e819 (2020).

17. Cele, S., et al. Escape of SARS-CoV-2 501Y.V2 variants from neutralization by convalescent plasma. medRxiv (2021).

18. Wang, P., et al. Increased Resistance of SARS-CoV-2 Variants B.1.351 and B.1.1.7 to Antibody Neutralization. bioRxiv (2021).

19. Choi, B., et al. Persistence and Evolution of SARS-CoV-2 in an Immunocompromised Host. N. Engl. J. Med. 383, 2291–2293 (2020).

20. Atyeo, C., et al. Distinct Early Serological Signatures Track with SARS-CoV-2 Survival. Immunity 53, 524–532 e524 (2020).

## References

21. Hadfield, J., et al. Nextstrain: real-time tracking of pathogen evolution. Bioinformatics 34, 4121–4123 (2018).

22. Walls, A.C., et al. Structure, Function, and Antigenicity of the SARS-CoV-2 Spike Glycoprotein. Cell 183, 1735 (2020).

23. Wang, Q., et al. Structural and Functional Basis of SARS-CoV-2 Entry by Using Human ACE2. Cell 181, 894–904 e899 (2020).

24. Berman, H.M., et al. The Protein Data Bank. Nucleic Acids Res. 28, 235–242 (2000).

25. McCallum, M., et al. N-terminal domain antigenic mapping reveals a site of vulnerability for SARS-CoV-2. bioRxiv (2021).

26. Cerutti, G., et al. Potent SARS-CoV-2 Neutralizing Antibodies Directed Against Spike N-Terminal Domain Target a Single Supersite. bioRxiv (2021).

27. Voss, W.N., et al. Prevalent, protective, and convergent IgG recognition of SARS-CoV-2 non-RBD spike epitopes in COVID-19 convalescent plasma. bioRxiv (2020).

28. Li, D., et al. The functions of SARS-CoV-2 neutralizing and infection-enhancing antibodies in vitro and in mice and nonhuman primates. bioRxiv (2021).

29. Suryadevara, N., et al. Neutralizing and protective human monoclonal antibodies recognizing the N-terminal domain of the SARS-CoV-2 spike protein. bioRxiv (2021).

30. Rapp, M., et al. Modular basis for potent SARS-CoV-2 neutralization by a prevalent VH1-2-derived antibody class. bioRxiv (2021).

31. Schoof, M., et al. An ultrapotent synthetic nanobody neutralizes SARS-CoV-2 by stabilizing inactive Spike. Science 370, 1473–1479 (2020).

32. Xiang, Y., et al. Versatile and multivalent nanobodies efficiently neutralize SARS-CoV-2. Science 370, 1479–1484 (2020).

33. Koenig, P.A., et al. Structure-guided multivalent nanobodies block SARS-CoV-2 infection and suppress mutational escape. Science (2021).

34. The PyMOL Molecular Graphics System, version 1.8 (Schrodinger, LLC, 2015).

35. Sholukh, A.M., et al. Evaluation of SARS-CoV-2 neutralization assays for antibody monitoring in natural infection and vaccine trials. medRxiv (2020).

